# Facilitating Electrochemical Lateral Flow Assay using NFC-Enabled Potentiostat and Nitrocellulose-based Metal Electrodes

**DOI:** 10.1101/2023.03.09.531916

**Authors:** Laura Gonzalez-Macia, Yunpeng Li, Kaijia Zhang, Estefania Nunez-Bajo, Giandrin Barandun, Yasin Cotur, Tarek Asfour, Selin Olenik, Philip Coatsworth, Jack Herrington, Firat Güder

## Abstract

Rapid detection of pathogens at the point-of-need is crucial for preventing the spread of human, animal and plant diseases which can have devastating consequences both on the lives and livelihood of billions of people. Colorimetric, lateral flow assays consisting of a nitrocellulose membrane, are the preferred format today for low-cost on-site detection of pathogens. This assay format has, however, historically suffered from poor analytical performance and is not compatible with digital technologies. In this work, we report the development of a new class of digital diagnostics platform for precision point-of-need testing. This new versatile platform consists of two important innovations: i) A wireless and batteryless, microcontroller-based, low-cost Near Field Communication (NFC)-enabled potentiostat that brings high performance electroanalytical techniques (cyclic voltammetry, chronoamperometry, square wave voltammetry) to the field. The NFC-potentiostat can be operated with a mobile app by minimally trained users; ii) A new approach for producing nitrocellulose membranes with integrated electrodes that facilitate high performance electrochemical detection at the point-of-need. We produced an integrated system housed in a 3D-printed phone case and demonstrated its used for the detection of Maize Mosaic Virus (MMV), a plant pathogen, as a proof-of-concept application.

## 1. Introduction

Rapid acquisition and communication of the results of a diagnostic test are essential for early detection and prevention of global pandemics impacting humans, animals or plants; there is, however, a lack of low-cost, versatile digital diagnostic platforms available in the market that can be easily adapted to unforeseen situations such as the rapid spread of known or unknown pathogens.

One of the most widely used Point-of-Need (PoN) tools for diagnostic testing and screening is the Lateral Flow Assay (LFA).^1–3^ LFAs are simple to use and do not require washing steps to run the assay. Conventional LFAs are performed using a strip, most commonly a nitrocellulose membrane, comprising a range of materials mounted on a backing card.^4,5^ Biorecognition and signal transduction are achieved using specific antibodies (or more recently their synthetic counterparts such as nucleic acid aptamers) tagged with colloidal gold or other color-generating particles.^6–8^ Results are interpreted based on the presence or absence of lines on the strip (*i.e*., nitrocellulose membrane), which can be read by the naked eye or with a reader. Most LFA devices, therefore, rely on colorimetric detection, which only provides qualitative or at best semi-quantitative results. Efforts have been made to create colorimetric LFAs that produce a quantitative output^9–11^ but the variability of light in the testing areas, presence of sample interferents (suspended solids, colored materials), and expensive instrumentation have hindered its proliferation.

An alternative strategy to obtain true quantitative outputs is the integration of electrochemical transduction into the LFA strips.^12,13^ Printing metal electrodes on nitrocellulose membranes using commercial inks has proven to be challenging however, due to the use of organic solvents in their formulations (needed to dissolve the polymer binder) which damage the delicate membrane. Instead of direct printing of metal inks onto the nitrocellulose membrane, attempts were made to attach external electrodes to the detection area of the LFA.^14–16^ Electrochemical measurement techniques naturally produce an electrical signal which can be easily digitized using low-power electrical circuits (compatible with miniaturization) and uploaded to the cloud over a smartphone. This is a critical consideration for the next-generation digital rapid diagnostics for on-site testing which can quickly inform individuals and decision makers to prevent the spread of pathogens. Despite the obvious advantages of electrochemical LFAs, tedious fabrication processes, complicated integration steps and the need for an external potentiostat to perform the measurements have prevented electrochemical LFAs from becoming a translational success.

One of the main challenges for digital rapid diagnostics is the need of a portable wireless reader for electroanalysis in the fields. Whitesides and coworkers reported a Bluetooth-based wireless potentiostat that can be controlled by a smartphone.^17^ Although high performance, the system required a battery for operation, could not be mounted on a smartphone and was unaffordable for deployment at scale in developing world. The use of additional batteries is particularly undesirable due to their impact on the environment.^18^ Bluetooth, WiFi and other longer-range wireless technologies may also be susceptible to breaches in data security, which is critical especially in health-related applications. Near-field communication (NFC) is a widely used wireless technology that enables secure transfer of data between devices at short distances (< 5 cm) using radio frequency signals at 13.56 MHz.^19^ The biggest differentiator of NFC, however, is that it is powered passively without batteries through wireless inductive coupling. NFC allows exchange of electrical power large enough (10 mW) to operate low-power and low-cost electronics (including microcontrollers and sensors).^20–24^ The elimination of batteries reduce cost, complexity and increase mobility since charging is no longer required.

In this work, we report two new key technologies to produce the next generation digital rapid diagnostics for on-site testing (**Figure 1**). We have developed: (i) a new approach for printing binder-free, hydrophilic 3D metal electrodes on/in nitrocellulose membranes to enable electrochemical LFAs; (ii) a programmable opensource wireless and batteryless NFC-enabled low-cost potentiostat to perform electrochemical measurements using a smartphone. The NFC-enabled diagnostics platform with <u>e</u>lectrochemical detection on <u>n</u>itro<u>c</u>ellulose membranes (eNC) can be easily adapted to lateral flow detection to achieve real-time identification of diseases in the field. We characterized the microstructural and electrochemical properties of the 3D electrodes for electroanalytical measurements. We also engineered the NFC-based potentiostat, manufactured a 3D-printed phone case and designed an Android application for use in the field. We demonstrated the operation of the integrated system (NFC-eNC) by measuring the product of a modified commercial enzyme-linked immunosorbent assay (ELISA) kit to detect Maize Mosaic Virus (MMV), a devastating crop pathogen. We compared the results of the electrochemical measurements to traditional spectroscopic ELISA measurements for validation of our approach for on-site testing.

**Figure 1.**
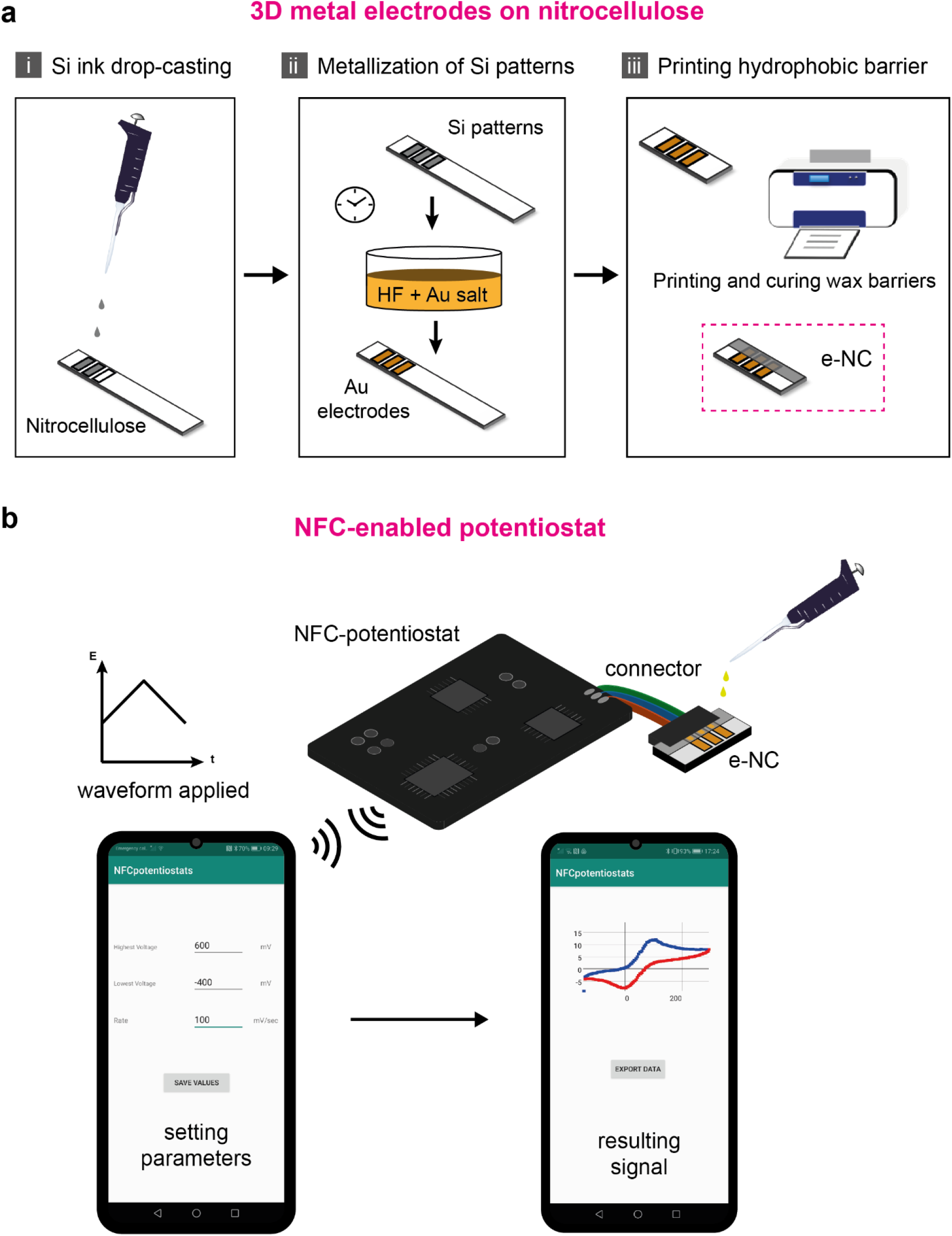
Schematic illustration of the key technologies developed in this work. **a** Fabrication of 3D metal electrodes on nitrocellulose membranes. **b** Batteryless, wireless NFC-enabled potentiostat to perform on-site electroanalytical measurements.

## 2. Results

### Design and operation of the NFC-enabled low-cost potentiostat

We designed an NFC-powered wireless potentiostat using an Arduino compatible low-cost and low-power 8-bit microcontroller (ATmega328P) to create an open-source development environment for applications in (chemical) sensing (**Figure 2A** and **Figures S1** and **S2**). Instead of an application specific integrated circuit (ASIC) to implement a potentiostat with a few pre-determined, fixed functions (such as the technology reported by Krorakai and coworkers and Silicon Craft Ltd.),^25^ our microcontroller-based approach is versatile and most electroanalytical measurement methods (such as cyclic voltammetry, chronoamperometry, *etc*.) can be implemented through software (**Figure 2B**). The microcontroller manages both the communication with the smartphone and the onboard potentiostat. The onboard, custom potentiostat consists of three operational amplifiers: U1 to U3. U1 is used to maintain the voltage set-point across the working (WE) and counter electrodes (CE) with respect to a third, reference, electrode (RE). U2 acts as a trans-impedance amplifier to convert the current collected at WE to a potential (relative to Voffset). U3 is used as a signal buffer to electrically isolate the sensor from the rest of the circuit. A dedicated 12-bit digital-to-analog converter (DAC; MCP47FEB22A0-E/S) defines the analog potential between WE and CE as well as the offset used to bias the sensed potential. An analog potentiometer (PVG3A104C01R00) scales the output range of the potentiostat to prevent clipping during signal digitization. Finally, a 10-bit analog-to-digital converter (ADC; part of Atmega328P) reads the resulting output of the potentiostat to enable further processing by the microcontroller and transmission to the smartphone **(Figure 2C** and **Table S1)**. For low-power operation, the microcontroller was clocked down to 1 MHz programmatically using the internal clock of ATmega328p which reduced power consumption of the microcontroller to ~300 μA at 1.8 V in active mode.

**Figure 2.**
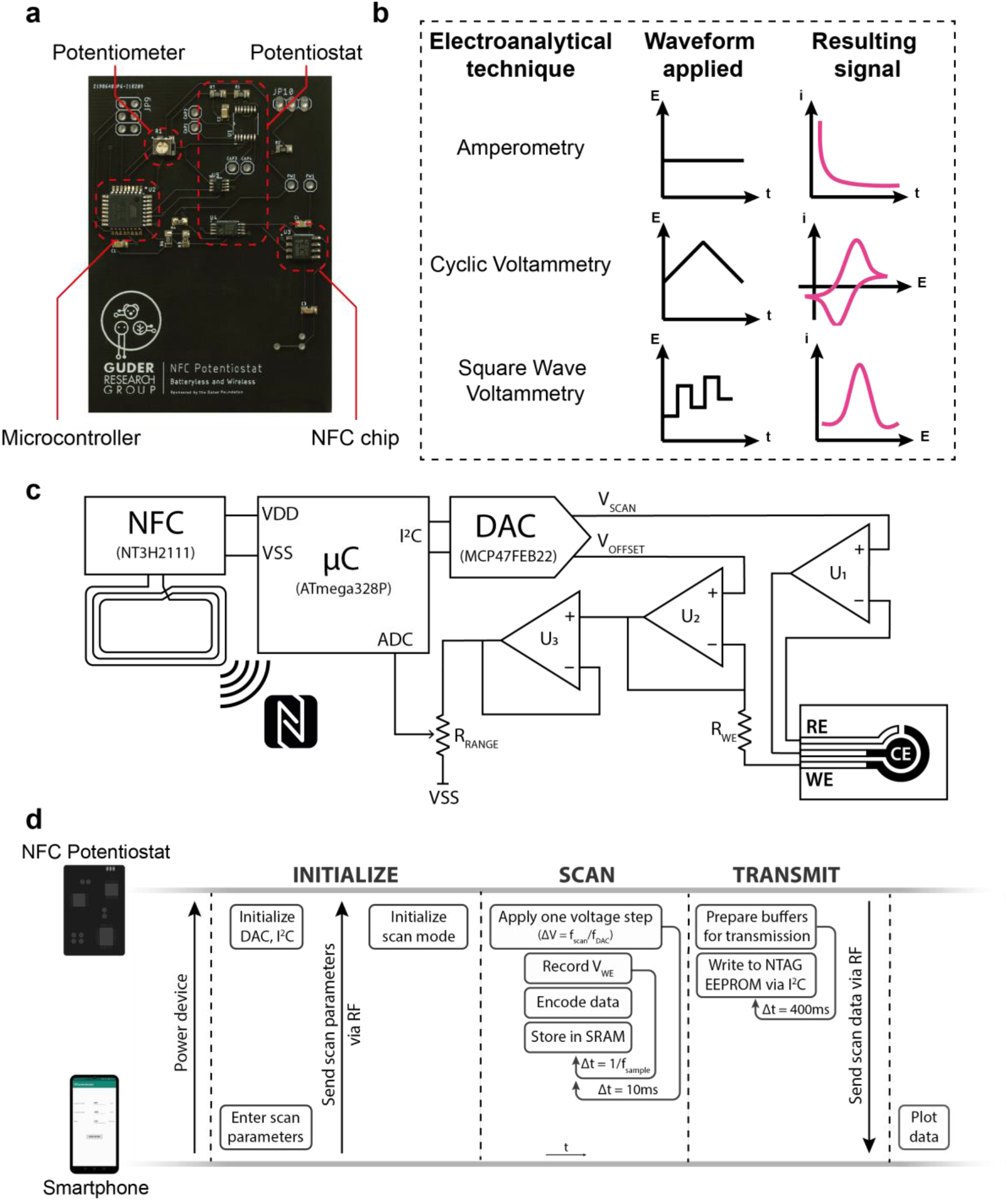
Operation and communication protocol of the NFC-potentiostat. **a** Photograph of the NFC-potentiostat. **b** Summary of the waveforms applied and resulting signals for three typical electroanalytical techniques performed with the NFC-potentiostat. **c** Scheme of the operation of the NFC-potentiostat: WE = working electrode, RE = reference electrode, CE = counter electrode, V_SCAN_ = scanning potential, V_REF_ = reference potential, R_POT_ = potentiometer, R_WE_ = resistor converting current through WE to voltage. **d** Communication protocol between the smartphone (app and NFC tag) and the NFC-potentiostat during the recording of a measurement: f_scan_ = scan rate, f_DAC_ = output rate of DAC, f_sample_ = sampling rate of potentiostat, t = time, ΔV = one voltage step, Δt = one time step, V_WE_ = WE potential.

The NFC-based potentiostat with all its components consumed less than 1 mA at 1.8 V (1.8 mW), nearly an order of magnitude below the nominal power budget for NFC-based power exchange (10 mW). Hence our design leaves more power budget to be used for future iterations by including additional components for new (analytical) functions or low-noise designs. The NFC-potentiostat can perform electrochemical measurements with a minimum range of potential between ± 0.9 V, which is sufficient for most electroanalytical experiments.

The NFC-potentiostat contains embedded software (written in C++) to communicate with external devices and run electroanalysis measurement protocols in coordination with a companion Android mobile application operated by the user. The communication between the NFC-potentiostat and the Android device is established through the 2 kilobyte (KB) non-volatile memory (EEPROM) as a buffer (**Figure 2D**). Using the Android mobile app, the user enters and saves the experimental parameters (**Figure S3**) to the memory of smartphone. Once the Android device is in sufficient proximity to the NFC-potentiostat, the smartphone wirelessly writes a command to the EEPROM of the NFC chip (NTAG) present on the NFC-potentiostat with the parameters of the electroanalytical measurement (Figure 2D). The embedded software on the NFC-potentiostat reads this command and runs the experiment, storing the results of the measurements (up to 400 datapoints - encoded as 800 bytes) on the on-board volatile 2 KB SRAM integrated within the ATmega328P chip. SRAM communication is chosen as the preferred medium for temporary data storage as the time taken to write directly to the NTAG’s EEPROM is significantly longer (<1 μs vs ~3 ms access time, respectively). When the measurement is completed, 400 samples (datapoints) collected in the SRAM buffer are transferred to the EEPROM of the NTAG and wirelessly communicated to the smartphone to share with the user. The results of the measurements can be visualized using the mobile app or shared with others over the cloud for further analysis.

### Electrochemical characterization of the NFC-potentiostat

Because our microcontroller-based design is highly flexible, a variety of 2- and 3-electrode electroanalytical methods can be programmatically implemented using the NFC-potentiostat. Due to their analytical importance, we have programmatically implemented cyclic voltammetry (CV), chronoamperometry (CA) and square wave voltammetry (SWV) which can be executed through function calls with the experimental parameters passed on as input arguments (see SI for source code and documentation). We compared the electroanalytical performance of the NFC-potentiostat to a commercial potentiostat (PalmSens4, PalmSens BV, The Netherlands) as shown in **Figure 3**. We used commercially available screen-printed carbon electrodes (SPCEs) and solutions of potassium ferrocyanide at concentrations ranging from 50 μM to 10 mM in 0.1 M KCl (**Figure S4**) for characterization. Fresh potassium ferrocyanide solution was placed on (all three) electrodes before each measurement. The electrodes were rinsed with deionized (DI) water and wiped dry between measurements. To perform a measurement, we first connected the screen-printed electrodes to the NFC-potentiostat (Figure 1B). Next, we selected the electroanalytical technique on the Android app with the experimental parameters. Finally, we added the sample solution onto the electrodes, and initiated the measurement by tapping the smartphone on the NFC-potentiostat. The smartphone remained in contact with the NFC-potentiostat during the measurement to ensure steady connection and exchange of power.

**Figure 3.**
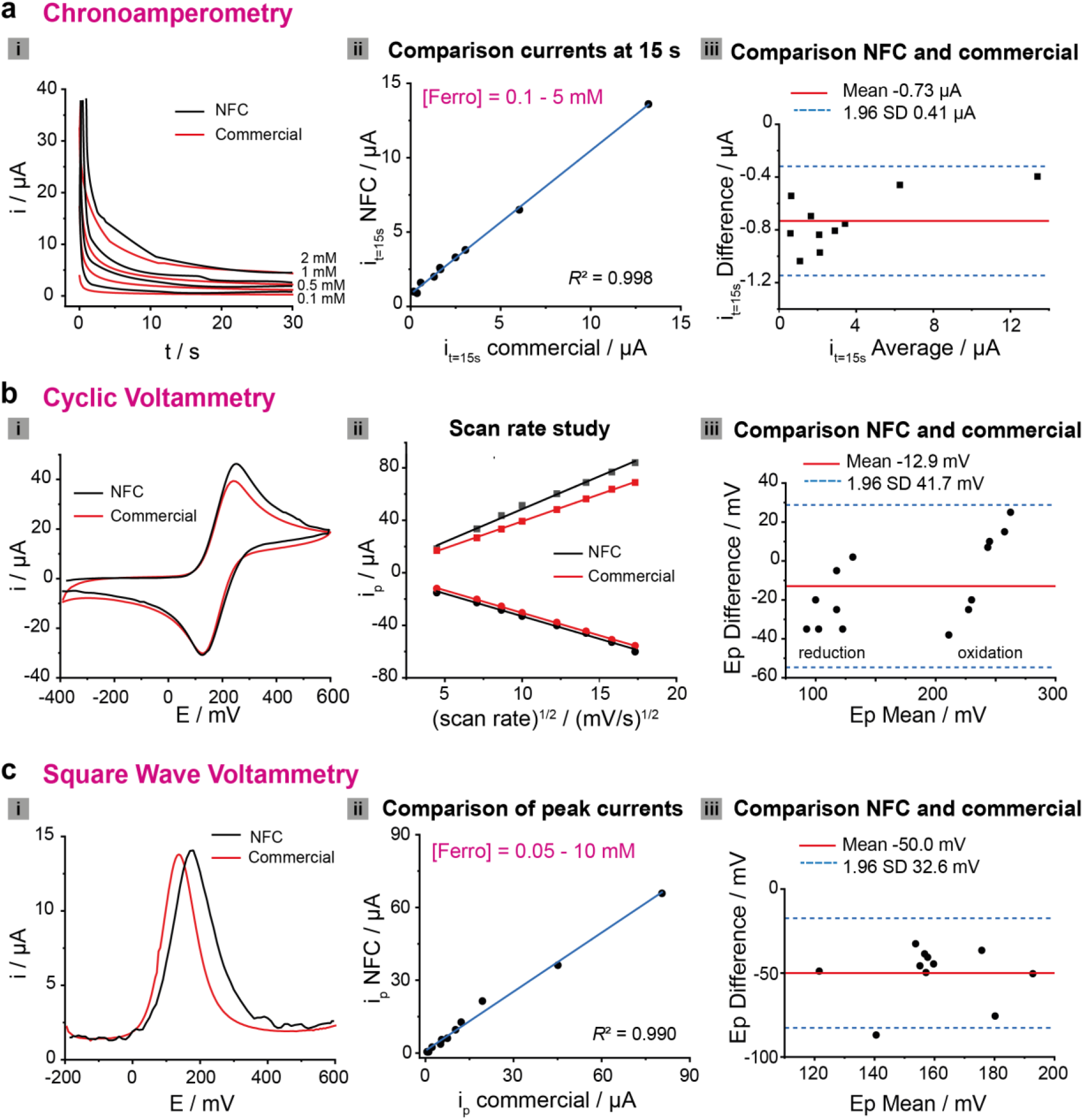
Electrochemical characterization of the NFC-potentiostat. Comparison NFC and commercial (PalmSens4) potentiostats using SPCEs: **a i.** Chronoamperometry at 200 mV (vs. Ag pseudo-reference) in several concentrations of potassium ferrocyanide; **ii**. Correlation of currents at t = 15 s measured by PalmSens potentiostat (commercial) and our NFC-potentiostat (ferrocyanide concentrations: 100 μM - 5 mM); **iii.** Bland-Altman plot of data in ii (SD, Standard Deviation). **b i.** Cyclic Voltammogram of 2 mM potassium ferrocyanide at 100 mV/s; **ii.** Scan rate study: linear correlation of peak current (i_p_) and square root of scan rates as expected for diffusion-controlled processes;^28–30^ **iii.** Comparison of commercial and our NFC-potentiostat by Bland-Altman plot of peak potentials (E_p_) from the scan rate study. **c i.** Square Wave Voltammetry of 1 mM potassium ferrocyanide (potential step: 5 mV; amplitude: 20 mV; frequency: 10 Hz); **ii.** Correlation of peak currents measured by commercial and our NFC-potentiostat using conditions as i (ferrocyanide concentrations: 50 μM - 10 mM); **iii.** Bland-Altman plot of peak potentials using conditions and ferrocyanide concentrations as ii.

#### Chronoamperometry (CA)

Chronoamperograms of solutions of potassium ferrocyanide at concentrations ranging from 50 μM to 10 mM were recorded using SPCEs at 200 mV (*vs*. Ag pseudo-reference) for 30 s (**Figure 3A**; for all chronoamperograms and the corresponding calibration curves please see **Figure S5**). Measurements performed with the NFC-potentiostat showed a good correlation with those obtained by the commercial potentiostat beyond 10 s, resulting in R^2^ = 0.998 (Figure 3A.ii). Bland-Altman plot (Figure 3A.iii) confirmed the good agreement between both potentiostats since all the values were within the limits of agreement (±1.96 SDS), with a consistent negative bias of the NFC-pontentiostat respect to the commercial one (Mean = - 0.73 μA).^26,27^ The difference in the initial currents (t < 10 s) could be corrected by pre-calibration of the instrument.

#### Cyclic Voltammetry (CV)

Cyclic voltammograms (CVs) of solutions of potassium ferrocyanide at concentrations between 100 μM - 5 mM were recorded at 100 mV/s with commercial SPCEs (**Figures 3B.i** and **S6**). The readings obtained by the NFC-potentiostat were comparable to those obtained with the commercial potentiostat (PalmSens4). Both instruments also produced similar peak shapes. At low concentrations of ferrocyanide (< 100 μM), the NFC-potentiostat produced no distinguishable peaks from the background; at concentrations higher than 5 mM, the NFC-potentiostat exceeded 100 μA (the limit for our platform which could only be extended through manual adjustment) and produced distorted shapes (data not shown). As expected for a diffusion-controlled electrochemical reaction, in 2 mM potassium ferrocyanide, peak currents were proportional to the square root of the scan rates (inset **Figure 3B.ii** and **Figure S7**).^28–30^ Peak currents and potentials recorded by both potentiostats were also similar, with higher non-faradaic currents in the CVs recorded with the NFC-potentiostat at scan rates > 150 mV/s. Bland-Altman analysis (Figure 3B.iii) confirmed a consistent bias of the NFC potentiostat respect to the commercial one (Mean = - 12.9 mV). Such difference of the NFC-potentiostat could be further addressed by pre-calibration of the system. Reproducibility of CV using the NFC-potentiostat was also evaluated by performing multiple scans in a 2 mM solution of potassium ferrocyanide at 100 mV/s (**Figure S8**). The peak currents of the repeats were within 5% Relative Standard Deviation (RSD) for both anodic and cathodic peaks in comparison to 1% RSD obtained with the commercial potentiostat.

#### Square Wave Voltammetry (SWV)

Square wave voltammograms of solutions of potassium ferrocyanide at concentrations ranging from 50 μM to 10 mM were obtained using SPCEs (**Figure 3C** and **S9**). Peak currents obtained with the NFC-potentiostat were comparable to those obtained with the commercial potentiostat as shown in Figure 3C.ii, resulting in R^2^ = 0.990. Bland-Altman plot of peak potentials (Figure 3C.iii) has a negative bias (mean = −50 mV), which means the peak potentials measured by our NFC-potentiostat were greater than those measured by the commercial potentiostat. Some variability in the peak shapes and potentials were observed with the NFC-potentiostat at low concentrations of potassium ferrocyanide due to lower faradaic currents generated at the electrodes. This can be addressed through pre-calibration of the instrument.

### Fabrication and characterization of Au electrodes on NC membranes

The fabrication of Au electrodes on NC membranes was performed using a two-step process developed previously by our group for metallizing paper and textiles:^31^ In step one, a water-based ink containing Si particles and carboxymethyl cellulose (acting as a stabilizer and viscosity modifier) is printed or drop-casted on the NC membrane in areas outlined with a fabric marker. In step two, the Si particles within the ink were autocatalytically transformed into metals by dipping into an aqueous bath containing metal ions and hydrofluoric acid (HF).^31^ This electroless approach produces binder-free, hydrophilic, 3D porous metal electrodes that are ideally suited for electroanalysis.

To create binder-free metal electrodes on NC, we first optimized the formulation of the Si-ink (including the particle size for Si and concentration of carboxymethyl cellulose, CMC, in the ink). The formulation of the ink was broadly optimized for commercial NC membranes with the pore size of 0.45 μm and 8 μm which are typically used in lateral flow assays.^5,32,33^ We evaluated two different sizes of Si particles for creating electrodes on NC: 100 nm and 3 μm. These average sizes for the Si particles were chosen because of the similarity in the pore size of the NC membranes. In addition to the optimization of the Si ink, we optimized the concentration of HF and Au salt for metallization of the Si particles. We used the resistance of the Au electrodes fabricated on NC membranes as the performance metric - lowest resistance would be the most optimum formulation and process. **Figure 4A** shows the optical micrographs of the Si particles (left) and Au electrodes (right) on NC membranes before and after the autocatalytic metallization process. Optimized fabrication conditions are summarized in **Figure 4B** and **Table S2**. The process optimized for NC membranes produced gold electrodes with a resistance below 10 Ω.cm^-1^ for the membranes with an average pore size of 0.45 μm and below 40 Ω.cm^-1^ for 8 μm (**Figure 4C**). The resistances obtained for our binder-free Au electrodes were similar to electrodes produced using commercial, binder containing metal inks.^34^ The resistance of our Au electrodes were significantly lower than commercially available carbon electrodes, however, which are the primary material in screen-printed electrochemical sensors used widely.^34^ It was observed that the combination of small Si particles (100 nm) with 0.45 μm NC membrane still led to very resistive Au electrodes, even at high concentrations of CMC (5 mg/mL) and Au salt (47 mM). This might be due to the high ratio particle-to-porous size, which would make the particles to penetrate the membrane and to spread rather than to stay on the surface to create a block pattern. At high concentrations of CMC, the ink viscosity was high, maintaining the Si particles in proximity within the membrane and leading to more conductive Au electrodes after the metallization process. Good Au electrode conductivities (< 40 Ω.cm^-1^) were obtained with 3 μm Si particles in both NC membranes at high concentrations of CMC (5 mg/mL for 0.45 μm and 12.5 mg/mL for 8 μm) and lower concentration of Au salt (24 mM) than in the first example. Following the protocol previously developed in our group, 5% HF was first used to assist the metallization process in the presence of Au ions.^35^ Further studies at different concentrations of HF (from 1 to 5%) showed that electrodes fabricated in 1% HF exhibited resistances in the same order of magnitude than those metallized in 5% HF, with improved electrochemical behavior (**Figure 4D**). 1% HF was, therefore, adopted as the optimized concentration for the autocatalytic metallization of gold electrodes on NC.

**Figure 4.**
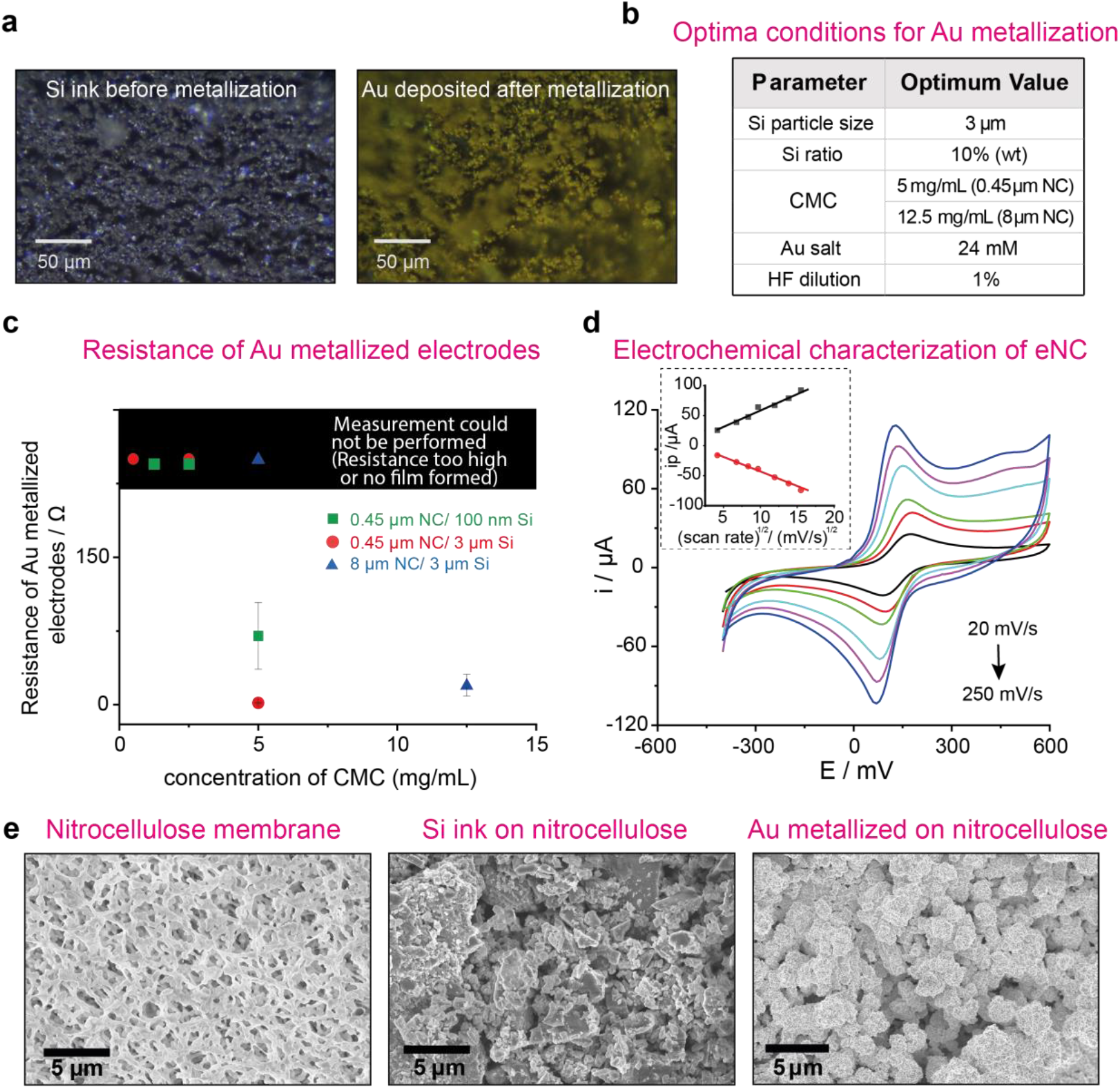
Fabrication and characterization of hydrophilic 3D metal electrodes on nitrocellulose membranes. **a** Optical micrographs of NC membranes with Si ink before metallization (left) and Au deposited after the autocatalytic metallization assisted by HF (right). **b** Optimized parameters for the fabrication of Au electrodes on NC membranes. **c** Resistances of Au electrodes after the metallization of Si particles depending on CMC concentration, NC porous size and Si particle size (n = 3) (Measurements could not be performed in cases where either Si ink solution overflowed the pattern barrier and strips were not good for further metallization or metallized electrodes peeled off during resistance measurements). **d** CV of 3D metallized Au electrodes in 2 mM potassium ferrocyanide/0.1 M KCl at scan rates from 20 to 250 mV/s; Inset: peak current *vs*. square root of scan rate 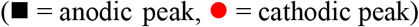. **e** SEM images (left to right: NC membrane, Si pattern electrodes, Au metallized electrodes).

Electrochemical characterization of the Au electrodes fabricated on NC was performed by cyclic voltammetry in 2 mM potassium ferrocyanide solution using a commercial potentiostat (Figure 4D). The middle electrode was used as WE, whereas the other two were used as RE and CE, respectively. Overall, peak separations were between 60 and 80 mV – higher than 57 mV expected for reversible one-electron redox reactions – indicating a quasi-reversible exchange between ferro/ferricyanide and the WE.^30^ Peak potentials were shifted approximately 50 mV towards zero potential in comparison with those obtained at the same scan rate and potassium ferrocyanide concentration using commercial screen-printed electrodes with Ag pseudo-reference electrode. Au electrodes have already been successfully used as pseudo-reference electrodes for sensing applications,^36^ and commercial electrodes are available in the market.^37^ Its potential range is, however, restricted in the presence of halide ions in solution.

Metallic gold suffers anodic dissolution and corrosion at oxidative potentials when ligands that form a stable complex with it are present in the electrolyte, losing their stability and suitability as WEs to study additional reactions.^38–40^ In this work, we use low oxidative potential range to avoid gold dissolution in the presence of chloride ions. Peak currents showed a linear relation to the square root of the scan rates (inset Figure 4D), as expected for diffusion-controlled electrochemical reaction.^28–30^ The effective areas of the Au WEs were calculated from the slopes (13.2 mm^2^ from the anodic data and 12.8 mm^2^ from cathodic data), similar to the geometrical area of 12 mm^2^ (3 mm x 4 mm).

Further characterization of the electrode surface was performed by Scanning Electron Microscopy. **Figure 4E** shows the electron micrographs of the NC membrane with a pore size of 0.45 μm before and after Si-ink deposition and following the Au-metallization process, recorded at 10.00K X magnification. The typical porous structure of the NC membrane observed on the first SEM image was fully covered by Si particles after the deposition of the precursor ink. The particles deposited on the NC membranes were of several sizes probably due to aggregation during the drop-casting and drying processes. After Au metallization, the NC membranes showed a particulate surface with higher homogeneously distributed structures. This was expected as a consequence of the autocatalytic metallization undergone on the Si surfaces in the presence of HF and Au^3+^ ions following Ostwald ripening process.^35,41,42^

### ELISA-based electrochemical detection of plant viruses

In agriculture, the surveillance of crops and early detection of plant diseases are crucial to avoid the spread of pathogens across geographies. To demonstrate the potential application of the technologies developed in this work for detecting plant pathogens, we applied the NFC-enabled digital diagnostic platform with eNC strips for detecting Maize Mosaic Virus (MMV).

Our detection approach is based on a commercial ELISA (enzyme-linked immunosorbent assay). Instead of relying on an optical output, however, we used the eNC strips to quantify the output of the assay products electrochemically with the future goal of creating a standalone test performed entirely using the LFA format in the field. The conjugated antibody utilized in the commercial Agdia ELISA Reagent set for MMV is labelled with the enzyme alkaline phosphatase (ALP), which catalyzes the hydrolysis of phosphate monoesters at high pH, forming inorganic phosphate.^43^ ALP is commonly used as a reporter in ELISA tests, using *para*-nitro phenyl phosphate (*p*-NPP) as a substrate (**Figure 5A**). The product of the enzymatic reaction, *para*-nitro phenol (*p*-NP), has a yellow color (**Figure 5B**) and its formation can be measured by spectrophotometry at 405 nm.^44^ The electrochemical detection of *p*-NP, however, requires the application of high overpotential resulting in an increase of interfering signals. *Para*-amino phenyl phosphate (*p*-APP) can also be used an alternative substrate for the ALP enzymatic reaction. The resulting product, *para* amino phenol (*p*-AP), however, unlike *p*-NP, is electroactive at lower potentials (**Figures S10** and **S11**), enabling electrochemical detection of the output of the assay with a higher performance.^45,46^ We performed the MVV ELISA using both enzymatic substrates, *p*-APP and *p*-NPP. The products of the test performed in a 96-well plate were measured by spectrophotometry at 405 nm after 1h from the addition of the Enzyme Conjugate substrate in 1 M DEA buffer pH 9.8 (Figure 5B). Subsequently, 75 μL of solution from each well was dropped onto commercial SPCEs and CV was performed at 100 mV/s. The electrochemical response of the ELISA products is shown in **Figure 5C**. The wells with positive control, where *p*-APP was used as the enzymatic substrate, exhibited the typical electrochemical response of *p*-AP, with an oxidation peak at 110 mV and reduction at 20 mV (*vs*. Ag pseudo reference). The measurements obtained were in agreement with the literature.^45,47^ The wells containing negative control and only GEB buffer with *p*-APP substrate and all the wells with *p*-NPP substrate did not show any electrochemical response, confirming the suitability of *p*-APP for the electrochemical detection of MMV virus following ELISA test.

**Figure 5.**
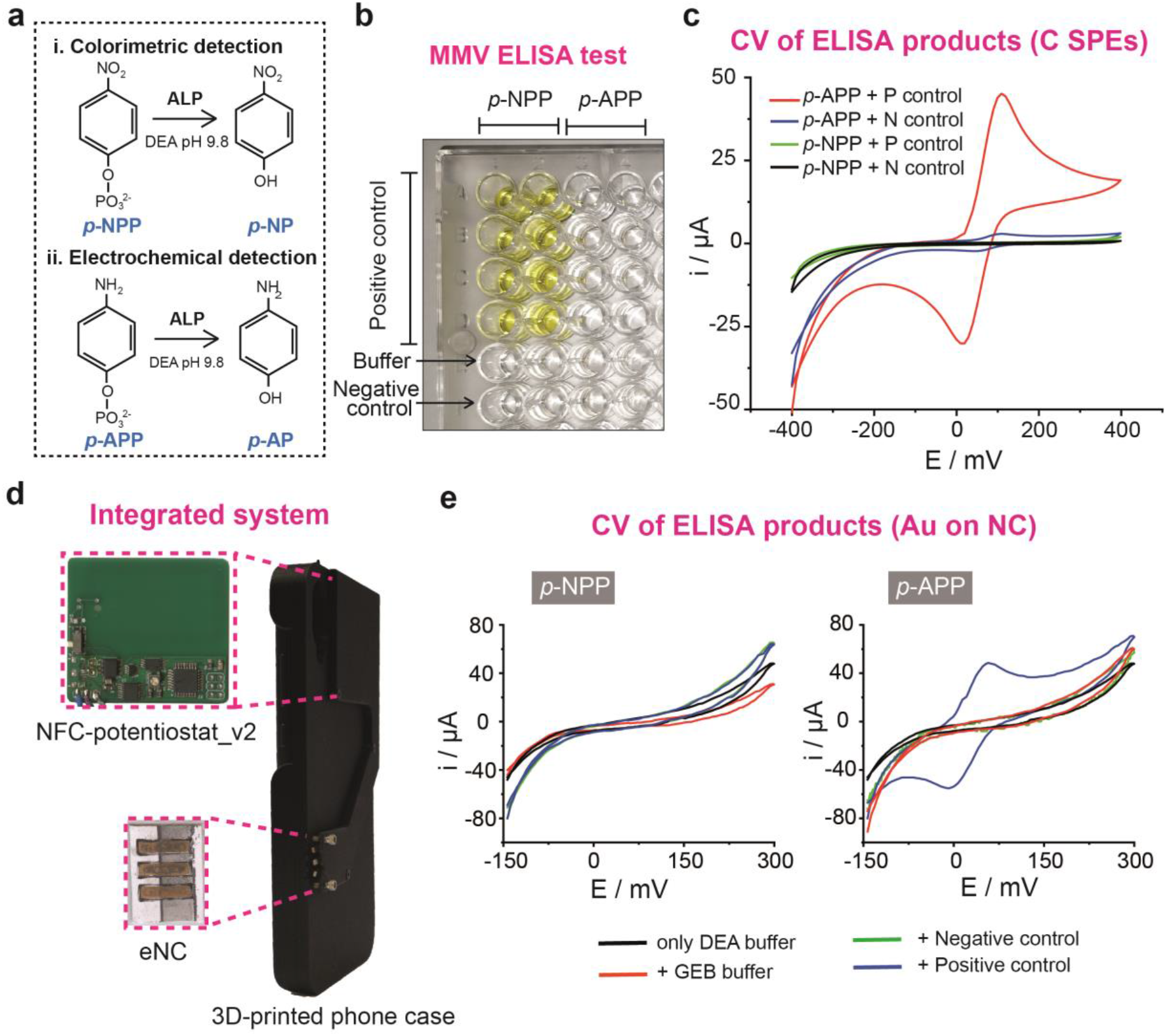
Spectrophotometric and electrochemical determination of the products of MMV ELISA test. **a** Scheme of enzymatic reactions of alkaline phosphatase (ALP) with *p*-NPP and *p*-APP as substrates. **b** Spectrophotometric determination of products obtained from the commercial MMV ELISA test using *p*-NPP and *p*-APP as substrates. **c** Electrochemical determination of same products as B, using commercial SPCEs and PalmSens4 potentiostat (P control = Positive Control; N control = Negative Control). **d** Picture of the integrated NFC-enabled diagnostics platform with NFC-potentiostat_v2 and eNC holder inserted into the 3D-printed phone case. **e** Cyclic Voltammograms recorded with the integrated system of ELISA products when *p*-NPP (left) or *p*-APP (right) were used as a substrate for the enzyme conjugate (WE area = 4.8 mm^2^). Cyclic Voltammograms have been smoothed using the Adjacent-Averaging method (10 points of window) in Origin 2021.

To ensure continuous communication during the measurements, a 3D-printed phone case was developed to house the NFC-potentiostat and connect the eNC strips (**Figure S12**). Housing the NFC-potentiostat in a phone case helped decrease the risk of loss in exchange of power and data during electroanalysis while providing the users with an integrated and easy-to-handle tool for crop diagnostics at the PoN. To integrate the NFC-potentiostat into the phone case, a smaller version was also fabricated (**Figure 5D**). The smaller NFC-potentiostat (NFC-potentiostat_v2, 4.2 cm x 4.7 cm) had the same circuit design as the first version (Figure 2 and S1) with the components compacted into an approximately 50% smaller area.

The eNC membranes with Au electrodes were first tested with the NFC-potentiostat and *p*-AP solutions (100 μM to 1 mM) to evaluate the system operation (**Figure S13**). The concentration range was chosen based on the CV peak currents of the ELISA products measured on SPCEs when *p*-APP and positive control were used in the assay (Figure 5C). Positive controls are used in commercial ELISA tests to ensure the assays run correctly and the output (Yes/No) response to the presence of the virus is reliable. Anodic peak currents of ELISA test products with positive control (1X) were approximately 45 μA, which was equivalent to the response of 600-650 μM *p*-AP (Figure S11). eNCs showed a linear response towards *p*-AP in 1 M DEA buffer pH 9.8 in the selected range of 100 μM to 1 mM. The plot of anodic peak currents *vs. p*-AP concentration is shown in the inset of Figure S13.

We finally applied the integrated system to measure the product of the MMV ELISA test on e-NC strips operated by the NFC-potentiostat and Android app (**Figures 5E**). MMV ELISA test was run with positive and negative controls and using both *p*-NPP and *p*-APP as substrates for the ALP enzymatic reaction. Cyclic voltammograms of the products were measured using our integrated system and the results collected by the app and stored in the cloud server. As expected, ELISA products after using *p*-NPP as enzymatic substrate did not show any electrochemical response in the range under study (Figure 5E left). When *p*-APP was used, ELISA products where positive control was added showed a well-defined electrochemical response, which allowed differentiation of positive controls from negative and blank samples (Figure 5E right).

## 3. Discussion

The NFC-eNC platform reported is highly versatile and can be modified to detect multiple analytes by tailoring the bioreceptors (i.e., antibodies, aptamers) or enzyme conjugates commonly used in LFAs. The NFC-potentiostat can be configured for simultaneous multiplexed testing by using its spare capacity for recording analog signals.

The NFC-potentiostat and 3D-printed phone case can be manufactured at a sufficiently low cost (US $18) for the laboratory prototypes. We expect the cost to be an order of magnitude lower at scale, produced using high volume industrial manufacturing technologies. Even for the laboratory prototypes, however, the cost of the NFC-potentiostat is below the price of commercial reduced-size potentiostats which typically sell for several hundred to thousands of USD. Because of its low-cost, the NFC-potentiostat can be employed both as a single use or reusable system to replace the conventional wired/wireless potentiostats commonly used in laboratories.

The integrated NFC-eNC system has at least three limitations: (i) We noted disagreements between the electroanalytical measurements recorded by the NFC-potentiostat and the commercial potentiostat at low currents. These disagreements in the recordings, however, can be addressed by pre-calibration of the NFC-potentiostat according to the range of currents of interest. (ii) The Si ink used for the deposition of the electrodes is manually drop-casted on NC membranes. The introduction of more automated techniques such as printers or low-volume dispensers can improve the reproducibility of this step and capacity of the process towards high-volume fabrication. (iii) The electrical connection between the eNC and phone case was at times unreliable. Nitrocellulose membranes are brittle, and electrical connector used for connecting the eNC to the NFC-potentiostat sometimes tear the membrane. Furthermore, in our current design, the eNC is exposed to the environment which can affect its performance. The connection of the eNC and the NFC-potentiostat can be improved by designing a custom eNC holder with fitted connections, where the eNC is protected from the environmental conditions.

With the adoption of LFA-based home testing due to the COVID-19 pandemic, most individuals have become accustomed to running rapid biological tests on their own at the PoN. The NFC-potentiostat and eNC membranes reported in this work aim to exploit this emerging trend and create the next generation of digitized tests for rapid detection of diseases at the PoN in agriculture, environment and healthcare.

## Methods

### Reagents

*Para*-nitrophenyl phosphate disodium salt (*p*-NPP), albumin from chicken egg powder, carboxymethyl cellulose sodium salt (CMC), diethanolamine (DEA) substrate buffer (5X) and nitrocellulose (NC) membranes (0.45 μm) were purchased from Thermo Scientific (UK). Nitrocellulose membranes (8 μm) were from Cytiva (formerly GE Healthcare Life Sciences, UK) and silicon metal powder (3 μm) was from Pilamec UK Ltd. *Para*-aminophenyl phosphate monosodium salt (*p*-APP), *para*-aminophenol (*p*-AP), *para*-nitrophenol (*p*-NP), bovine serum albumin (BSA), gold (III) chloride trihydrate, hydrofluoric acid (50%), magnesium chloride, phosphate buffered saline, polyvinylpyrrolidone, potassium chloride, potassium hexacyanoferrate (II), silicon nanopowder (< 100 nm), sodium bicarbonate, sodium citrate tribasic, sodium carbonate, sodium sulfite and TWEEN® 20 were obtained from Sigma-Aldrich, UK.

ELISA Reagent set for Maize Mosaic Virus (MMV) and Positive Control for MMV were purchased from Agdia EMEA (Soisy-sur-Seine, France). DropSens screen printed carbon electrodes (SPCEs) were purchased from Metrohm UK Ltd (Runcorn, UK). They are comprised of a carbon working electrode (4 mm diameter), a carbon counter electrode and a silver reference electrode (**Figure S5**). Wax ink (black, yellow, cyan, and magenta) for Xerox ColorQube 8580 printer were purchased from local Xerox distributor.

Materials for the 3D-printed phone case were sourced from EnvisionTec and Raise3D.

### NFC-potentiostat design and fabrication

The reusable NFC-potentiostat comprises a printed circuit board (PCB) and Android app for data visualization and communication with a cloud server. The printed circuit board (PCB) was designed using EAGLE by Autodesk Inc to perform data acquisition and wireless transmission to a mobile device through NFC communication. An NFC chip (NXP NT3H2211) provided by Digikey is used to harvest energy from and communicate with the user’s smartphone. The PCB (5 cm x 7 cm) was manufactured by Elecrow Ltd. and assembled in our lab. The Atmega328P, a widely used low-power programmable microcontroller, is integrated to setup and execute electrochemical scans, process incoming sensor data and handle all data communication. The operation of the NFC-potentiostat with the Android app is simple: the scan parameters of the experiment are entered in the app which, upon contact with a smartphone, are transmitted to the NFC-potentiostat to initiate electroanalysis of the sample. The current generated at the counter electrode is measured by the ADC of the microcontroller using a current-to-voltage converter. An analog potentiometer is embedded between the potentiostat and ADC to improve signal measurement (e.g., scaling up to increase signal resolution or scaling down to prevent clipping). The mobile application was developed using the Java programming language in Android Studio and then uploaded to a Huawei Lite P20 smartphone. The NFC antenna in the smartphone and the designed circuit board should be close in proximity to ensure proper transmission of power and data among the two units, enabling electroanalytical methods including: cyclic voltammetry, square wave voltammetry and chronoamperometry. The sampling rate of the collected signals is set to 10 Hz. The firmware is designed to send a maximum of 400 data points per experiment from the device to the smartphone: this allows for a scan of maximum 40 s duration. Scans of longer duration will require down-sampling of the sampling rate. The data is plotted after the full cycle has completed with an additional 10 s delay.

### Mobile phone application

The Android app developed is designed to run cyclic voltammetry (CV), chronoamperometry (CA) and square wave voltammetry (SWV), three of the most used electrochemical techniques for sensing applications. The NFC-potentiostat receives the parameters (i.e., scan type, scan rate, signal amplitude and scan range) and harvests power from the smartphone’s NFC functionality to conduct electrochemical measurements, providing a quick response in up to 40 s (depending on the scan rate). After data collection, the resulting signal is plotted graphically for visual inspection and ready to export. Android’s inherent data sharing functionality is used to allow for data sharing via email, Gmail and Bluetooth. The app also permits the transfer of data measured to cloud services such as Google Drive. For ordinary end-users, a simplified version of the Android app with pre-set parameters for electrochemical testing can be used. Results can then be provided using simple symbols to indicate the presence/absence of the target analyte.

### Fabrication of Au electrodes on NC membranes

The process to fabricate 3D metal electrodes on NC membranes comprised four steps: (i) Transfer of the 3-electrode design to the NC membranes, using a permanent fabric marker and laser-cut Cu mask; (ii) Drop-casting of the Si precursor ink; (iii) Metallization of the silicon patterns into Au, assisted by HF solution; (iv) Fabrication of hydrophobic barriers by wax printing to separate the electrode contacts from the detection area.^48,49^ The Si precursor ink consisted of Si particles (1 g), carboxymethyl cellulose (CMC) and DI water (10 mL), based on previous ink formulations developed in our group. The Si-ink patterns deposited on the NC membranes were then autocatalytically metallized in a bath containing AuCl3 in a dilute aqueous solution of hydrofluoric acid (HF).^35^ 1 00-nm and 3-μm Si particles were tested here, with CMC acting both as a stabilizer and viscosity modifier. Optical microscope images were taken on a Brunel SP202XM metallurgical microscope connected to a Nikon D3200 camera. Scanning Electron Microscopy (SEM) images of Au electrodes fabricated on NC membranes were acquired using a Zeiss Gemini Sigma 300 FEG electron microscope at 5 keV electron beam energy, unless otherwise specified.

After the electrode fabrication, a hydrophobic wax barrier was deposited perpendicular to the electrodes, along the strip. The barrier was first printed on a plastic substrate using a regular office wax printer (Xerox ColorQube 8580) and then transfer to the NC membrane by heat press (Vevor HP230B heat press) at 100 °C, creating a 5 mm wide LFA channel. The modified NC membranes were attached to a plastic substrate using double-sided tape to facilitate the handling and further measurement of the electrochemical LFA strips.

### ELISA

Agdia commercial ELISA kit for Maize Mosaic Virus (MMV) is based on Double Antibody Sandwich (DAS) assays using polyclonal antibodies and alkaline phosphatase (ALP) as the test label.^50^ Briefly, 96-well plates were incubated overnight in the fridge with the provided capture antibody (CAB) at 1:200 v/v dilution in coating buffer. After washing three times, 100 μL of positive control, negative control and general extract buffer (GEB) normally used to prepare real samples were added to the wells and the plate was incubated for 2 h at room temperature. Following eight washing steps, 100 μL of the enzyme conjugate antibody (ECA) at 1/200 v/v dilution in ECI buffer were added and the plate was incubated for other 2 h at room temperature. Finally, the plate was washed 8 times and 100 μL of enzyme substrate was added to the wells with some modifications from the manufacturer’s protocol (see section *ELISA-based electrochemical detection of plant viruses*). After 1 h incubation at room temperature, the absorbance of the plate at 405 nm was measured using a Varioskan microplate reader (Thermo Scientific) and SkanIt Software (version 2.4.5.9) Research Edition. The plates were incubated in an airtight container with a wet paper towel and the washing steps were performed with phosphate buffered saline solution containing Tween 20 (PBST, Agdia User Guide Buffer formulations).^50^ After the absorbance measurement, the solution of each well was transferred to 500-μL Eppendorf microtubes and subsequently measured electrochemically.

### Fabrication of 3D-printed phone case

The phone case was designed for a Huawei P20 Lite smartphone used in the measurements. The case consisted of two parts, with the spring-loaded contacts for the electrodes and the electronics board sandwiched in between, secured using M2X8mm and M2×6mm hex countersunk screws. The 3D-printed phone case was fabricated using a Prusa MK3S+ printer and polylactic acid plastic filament. The design was first made on Autodesk Fusion 360 software and sliced using PrusaSlicer software. The average print time was 8 h but it can be reduced to 3 h if printed using a stereolithographic printer.

## Data availability

The datasets generated during the current study are available from the corresponding author on reasonable request.

## Acknowledgements

The authors would like to thank the department of Bioengineering at Imperial College London. F.G. and L.G.-M. thank the Bill and Melinda Gates Foundation (Grand Challenges Explorations scheme under grant numbers OPP1212574 and INV-038695) and the European Union’s Horizon 2020 research and innovation program under the Marie Sklodowska-Curie grant agreement No 101025390. F.G. and E.N.-B. would like to thank the Wellcome Trust (Grant No. 207687/Z/17/Z). Y. C. would like to thank the Turkish Ministry of Education, EPSRC IAA and Innovate UK (10027758). F.G., J.H. and P.C. thank the EPSRC (EP/L016702/1). S.O. acknowledges the Imperial President’s PhD Scholarships. F.G., J.H. and P.C. would like to acknowledge Imperial College Centre for Processable Electronics and the Centre for Doctoral Training in Plastic Electronics.

## Author Contributions

These authors contributed equally: Laura Gonzalez-Macia, Yunpeng Li.

## Competing Interests statement

The authors declare no competing interests.

## Notes

### Competing Interest Statement

The authors have declared no competing interest.

